# Interaction of habitat isolation and seasonality: impact of fragmentation in dynamic landscapes on local and regional population variability in meta-food chains

**DOI:** 10.1101/2020.09.16.299818

**Authors:** Markus Stark, Moritz Bach, Christian Guill

## Abstract

While habitat loss is a known key driver of biodiversity decline, the effect of habitat fragmentation ‘per se’ is far less clear. Environmental perturbations, e.g. due to seasonality, potentially interact with effects of habitat fragmentation, thus making the latter even more difficult to assess. In order to analyze how the dynamic stability of local and metapopulations is affected by habitat fragmentation, we simulate a meta-food chain with three species on complex networks of habitat patches and evaluate the average variability of the local populations and the metapopulations as well as the level of synchronization between the patches. To account for environmental perturbations, we contrast simulations of static landscapes with simulations of dynamic landscapes, where 30 percent of the patches periodically become unavailable as habitats. We find that it depends on the parameterization of the food chain whether higher dispersal rates in less fragmented landscapes synchronize the dynamics or not. Even if the dynamics synchronize, local population variability can decrease due to indirect effects of dispersal mortality, thereby stabilizing metapopulation dynamics and reducing the risk of extinction. In dynamic landscapes, periodic external perturbations often fully synchronize the dynamics even if they act on a much slower time scale than the ecological interactions. Furthermore, they not only increase the variability of local and metapopulations, but also mostly overrule the effects of habitat fragmentation. This may explain why in nature effects of habitat fragmentation are often small and inconclusive.

## 1 Introduction

Anthropogenic habitat degradation and loss are strong negative drivers of biodiversity on local and global scales (Butchart et al., 2010, Pereira et al., 2010, Pimm et al., 2014). While habitat loss has a clear cause-effect relationship with declining diversity induced by e.g. lack of resources, habitat size restrictions, or increased mortality (Brooks et al., 2002, Duraiappah et al., 2005), the effect of habitat fragmentation ‘per se’ is still intensely debated (Fahrig, 2017, Fahrig et al., 2019, Fletcher et al., 2018, Hanski, 2015). Following Fahrig (2003), habitat fragmentation comprises attributes of spatial configuration and the size of habitat patches, but excludes habitat loss. Effects of fragmentation ‘per se’ are more difficult to assess because they are usually weaker than the effects of habitat loss (Fahrig, 2003) and often confounded with the latter (Didham et al., 2012). Moreover, empirical and experimental fragmentation studies report conflicting results: Negative effects on diversity have been attributed to the prevention of rescue effects (Gotelli, 1991, Levins, 1969), positive effects can arise due to an increase of local diversity in more isolated habitats (Fahrig, 2017), and increases of habitat edges in more fragmented landscapes can have combinations of positive and negative effects (Pfeifer et al., 2017).

Theoretical models allow to disentangle the effects of habitat loss and habitat fragmentation more easily. Classical colonisation-extinction models demonstrated, for example, that habitat destruction may favour species that are inferior in competition, but superior dispersers (Nee and May, 1992, Tilman et al., 1994) or how deconstruction of food webs in response to habitat loss depends on the structure of the web (Melián and Bascompte, 2002, Pillai et al., 2011). However, these models only take into account presence or absence of a species on a patch but not the population dynamics within a patch.

A major concern of models that include explicit population dynamics are mechanisms that synchronize population cycles between habitat patches. Such synchronous oscillations destabilize metapopulations by amplifying the amplitude of oscillations in their regional abundances and increasing the extinction risk of species in entire regions due to correlated local extinction events. Conversely, asynchronous oscillations can promote regional persistence and stability through rescue effects (Blasius et al., 1999, Levins, 1969) or averaging (Briggs and Hoopes, 2004). These models, which are often limited to either a small number of patches or to regular, rectangular lattices (Briggs and Hoopes, 2004), have established that the synchronicity of population oscillations between patches generally increases with dispersal rate (Jansen, 2001, Sherratt et al., 2000). Other factors affecting synchronicity are adaptive dispersal (Abrams, 2007, Abrams and Ruokolainen, 2011), inter- and intraspecific density dependence of dispersal rates (Hauzy et al., 2010), and costliness or distance dependence of dispersal (Koelle and Vandermeer, 2005). In larger networks of habitat patches, an irregular network structure favours asynchronous dynamics (Holland and Hastings, 2008), but high dispersal rates again lead to synchronous oscillations that are detrimental for species persistence (Plitzko and Drossel, 2015).

The dispersal rate is directly related to how well patches are connected to each other or how large the effective distance between them is (Fletcher et al., 2016, Koelle and Vandermeer, 2005). As a more fragmented landscape leads to a decoupling of habitats and limits landscape connectance, this connects results regarding synchronisation of population oscillations with research on the effect of habitat fragmentation. Indeed, it has been shown that synchronization among natural populations declines with increasing distance between them (Ranta et al., 1995).

Another major driver of synchronized population oscillations are correlated environmental fluctuations (Kahilainen et al., 2018, Koenig, 1999, Moran, 1953, Ranta et al., 1995). These fluctu-ations can affect demographic rates of the species via changing environmental conditions (like ambient temperature or resource availability), but they can also directly influence the availability of patches as habitable areas. As an example, a landscape in which both a temporally variable environment and a pronounced spatial structure strongly affect ecological communities are kettle holes in formerly glaciated regions (Kalettka and Rudat, 2006). These small ponds are typically formed in large clusters, and seasonal changes of temperature and precipitation cause some of them to be only temporally filled with water. The local aquatic communities of these temporary ponds thus regularly become completely extinct and, besides resting stages, recolonisation through dispersing species from permanent ponds is a key element to reestablish the communities and preserve local diversity (De Meester et al., 2005). A successful recolonization hereby depends on the spatial structure of the landscape and individual species attributes like the dispersal strategy (Vanschoenwinkel et al., 2009). Long distances between habitats in an otherwise inhospitable landscape can cause a lower connectance and increase travel costs, which decrease the dispersal success of a species.

This means that successful dispersal and hence the preservation of diversity can depend on both temporal and spatial attributes of a landscape. However, few studies specifically address the interplay between environmental fluctuations and habitat fragmentation (but see (Gouhier et al., 2010) for an example of interaction between dispersal rate and correlated environmental fluctuations).

Here we study the dynamics of a tri-trophic meta-food chain on a large, spatially explicit network of habitat patches and analyze its stability with respect to (1) habitat fragmentation (measured by the connectance of the network of habitat patches) and (2) seasonal environmental disturbances of the landscape (dynamic landscape; periodic extinction of a subset of local communities when their patches are temporarily unavailable as habitat). In order to obtain a complete picture of the effects of habitat fragmentation and seasonal environmental disturbances on the extent and synchronicity of population oscillations in a food chain, we analyse two parameterizations of the food chain that correspond to contrasting oscillation patterns. These patterns are characterised either by a relatively even distribution of biomass along the food chain (weak trophic cascade) or by marked differences among the species (strong trophic cascade), both of which are common in natural ecosystems (Carter and Rypstra, 1995, Estes and Duggins, 1995).

Our model setup explicitly addresses one aspect of fragmentation, namely the connectance of habitats, and omits the effects of habitat loss. We consider both static landscapes, where all patches are constantly available as habitats, and dynamic landscapes, where seasonal environmental fluctuations periodically render some of the patches unavailable as habitats. The stability of the dynamics of the metacommunity is evaluated within the framework of Wang and Loreau (2014) that divides population variability into an *α*-, *β*-, and *γ*-component (similar to the classical diversity indices by Whittaker (1972)): *α*-variability is the average coefficient of variation of a species’ local abundances, *γ*-variability is the coefficient of variation of the regional (metapopulation) abundance, and *β*-variability quantifies differences in oscillations between patches, i.e., how synchronous the local populations oscillate. We expect that increasing habitat connectance (i.e., decreasing fragmentation) increases dispersal flows, as it opens up more opportunities for individuals to move between patches and decreases mortality during dispersal. This will synchronise population oscillations, which will be further enhanced by seasonal (synchronous) environmental fluctuations. Furthermore, we expect local (*α*-) variability to increase, as declining dispersal mortality allows more biomass to flow up the food chain, thus strengthening (and thereby destabilising) the trophic interactions (Rip and McCann, 2011). If the local population oscillations indeed become more synchronous, this will also increase regional (*γ*-) variability as habitats become more connected.

## 2 Methods

The model comprises a tri-trophic food chain including an autotroph (*A*), a consumer (*C*) and a predator (*P*) species. As basis for the growth of the autotroph a dynamic resource (*R*) serves as essential energy source and can be seen as a universal nutrient. This food chain is extended to a metacommunity by placing copies of it on habitat patches that are randomly distributed in space and connected via species-specific dispersal links (Fig. 1).

**Fig. 1.**
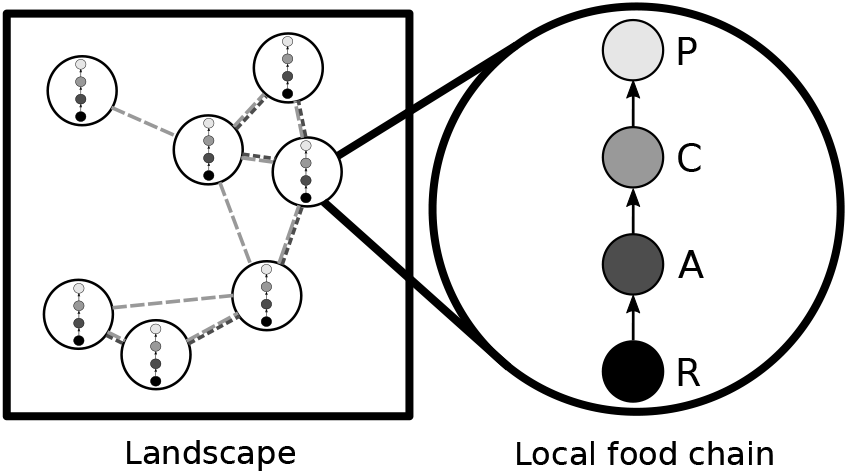
Left: simplified example of a spatial network of habitat patches. Dashed lines of different grey tones indicate dispersal links of the respective species. The resource does not disperse between patches. Right: local food chain on each patch comprising three trophic levels (autotrophs, ***A***, consumers, ***C***, and predators, ***P***) plus a dynamic resource, ***R***.

### 2.1 Trophic interactions

We first describe only the trophic interactions between the populations on a single patch and disregard dispersal. The local dynamics of the food chain follow a generalization of the bioenergetics approach (Brose et al., 2006b, Yodzis and Innes, 1992), supplemented with an equation for the resources. Adapted from chemostat dynamics, the rate of change of the resource density *R* is expressed as

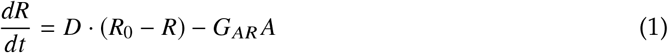

with the resource turnover rate *D* and the supply concentration *R*_0_. Uptake of resources by the autotroph *A* is described by a Monod function 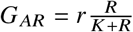 with maximum uptake rate *r* and half saturation constant *K*. The rates of change in biomass density for each species (*A, C* and *P*) are expressed by

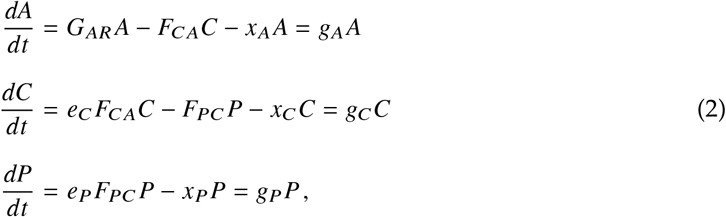

where the first terms in all three equations represent growth due to consumption, the last terms denote metabolic losses, and the middle terms in the equations for the autotroph and the consumer describe mortality through predation. The terms are summarized by the net per capita growth rates *g_i_*(*i* = *A, C, P*). The parameters *e_i_* and *x_i_* are assimilation efficiencies and per capita respiration rates, respectively. The per capita feeding rate of species *i* on species *j* is described by a Beddington-DeAngelis functional response (Beddington, 1975, DeAngelis et al., 1975):

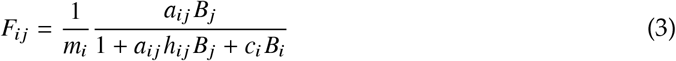

with the attack rate *a_ij_*, the handling time *h_ij_*, the interference coefficient *c_i_*, and *B_i_* and *B_j_* as placeholders for the respective consumer’s or resource’s biomass density. Since the model is formulated in terms of biomass densities (as opposed to population densities), the functional response is scaled with 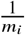, the inverse of the respective consumer’s body mass (Heckmann et al., 2012).

The parameters of the trophic dynamics scale allometrically with the body mass of the species. Mass-specific maximimum growth rate and respiration rates are assumed to decrease with a negative quarter-power law with body mass, i.e. 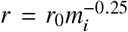 and 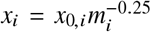 (Brose et al., 2006b, Yodzis and Innes, 1992). Following Rall et al. (2012), handling times depend on the body masses of both consumer and resource with 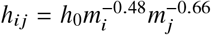. The same is true for the attack rates, but since these parameters were used to differentiate the contrasting states of top-down control, fixed values were used here (c.f. Tab. 1) that nevertheless obey the general trends found in (Rall et al., 2012). Body masses increase by a factor of 100 per trophic level, a value commonly found in invertebrate communities and known to have a stabilising effect on population dynamics (Brose et al., 2006a,b). Freedom of choosing an appropriate set of units allows us to set the body mass of the autotroph to *m_A_* = 1. In general, the model is parameterized such that the population dynamics of all species are oscillatory when dispersal is not accounted for (Tab. 1, Fig. 2).

**Fig. 2.**
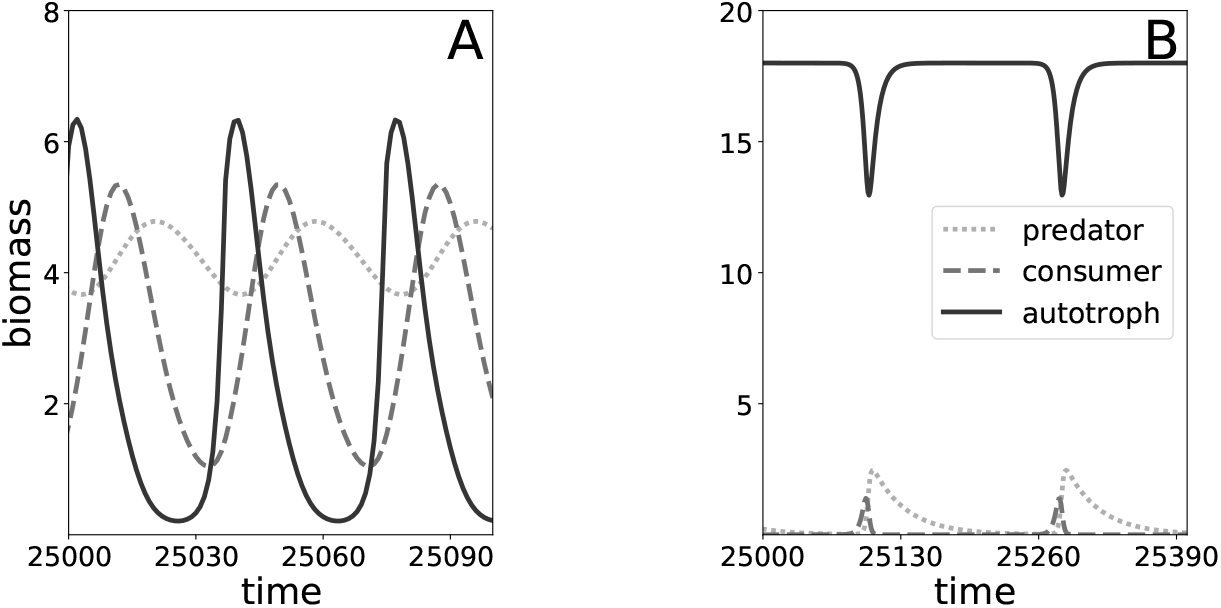
Timeseries of the dynamics for the weak (A) and the strong (B) trophic cascade on a single patch without dispersal dynamics. In case A, *a_CA_* = 105 and *a_PC_* = 450; in case B, *a_CA_* = 170 and *a_PC_* = 10000. All other parameters as in Tab. 1. Note the different scales of x- and y-axes in the two panels.

**Table 1.** Standard parameter set used in the model.

| Parameter | Description | Value |
| --- | --- | --- |
| $D$ | resource turnover rate | 0.5 |
| $R_0$ | resource supply concentration | 5 |
| $r_0$ | intercept mass specific max. resource uptake rate | 1 |
| $K$ | half saturation density for resource uptake | 0.2 |
| $c_C, c_P$ | interference competition | 0.6 |
| $e_C$ | assimilation efficiency consumer ($C$) | 0.45 |
| $e_P$ | assimilation efficiency predator ($P$) | 0.85 |
| $x_{0,A}$ | intercept respiration rate plant ($A$) | 0.138 |
| $x_{0,C}, x_{0,P}$ | intercept respiration consumer ($C$) and predator ($P$) | 0.314 |
| $a_{AC}$ | attack rate consumer | 105 or 170 |
| $a_{PC}$ | attack rate predator | 450 or 10000 |
| $h_0$ | intercept handling time | 0.1 |
| $D_0$ | intercept maximum dispersal distance | [0.06: 0.5] |
| $\epsilon$ | scaling exponent for maximum dispersal distance | 0.05 |
| $\mu_0$ | scaling factor maximal emigration rate | 2 |
| $b$ | curvature of emigration function | 25 |
| $Z$ | number of habitat patches | 30 |
| $\sigma$ | fraction of habitat patches blinking | 0.3 |
| $\lambda$ | period length of blinking cycle | 6000 |

### 2.2 Habitat network and dispersal

We use the same rules for modeling spatial interactions as in Ryser et al. (2019). Dispersal is considered for the autotroph, consumer, and predator species in the model. The spatial setting is implemented as a random geometric graph (RGG) (Penrose, 2003), where each node of the spatial network represents a habitat patch for a local community (Urban and Keitt, 2001). The (x, y)-coordinates of each patch were drawn at random from a bivariate uniform distribution over the intervall [0: 1] × [0: 1]. Dispersal links between the patches connect the local populations, enabling exchange of biomass between patches and thereby forming a meta-food chain (Fig. 1).

Each species perceives its individual dispersal network depending on its body mass *m_i_*. A dispersal link for species *i* exists between two patches *k* and *l* only if the distance between them is less than the species-specific maximum dispersal distance

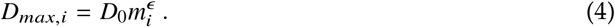

The exponent *ϵ* is set to a positive value to account for increased mobility and thus improved dispersal abilities of species with a larger body mass (Hein et al., 2012, Peters, 1983).

Dispersal itself is at least for animal species often an active process resulting in metabolic costs and potentially involving a higher risk of predation. To account for these costs, we assume that dispersal success *S_i,lk_* of species *i*, when moving between patches *l* and *k*, decreases linearly with the distance between the patches:

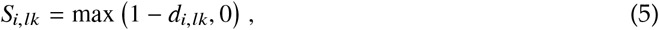

where 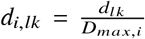 is the distance between the patches relative to the maximum dispersal distance of species *i*. For passively dispersing plants distance-depending costs can be caused by a decreasing probability of propagules finding by chance a suitable patch that is further away.

The fraction of individuals emigrating from a source patch *k* that move towards a target patch *l* is calculated using the weight function

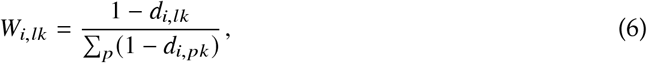

where the sum in the denominator is taken over all potential target patches *p* that are within the maximum dispersal range of species *i* on patch *k* (i.e., those with *d_pk_* < *D_max,i_*). This weight function makes dispersal links between nearby patches stronger than those between patches that are further apart. Note that while specific distances *d_i,lk_* and success terms *S_i,lk_* are symmetric for all pairs of patches, the weight function is not (i.e. *W_i,lk_* ≠ *W_i,kl_*).

In general, the process of dispersal can be described as an exchange of biomass between habitat patches that is affecting the population dynamics of species *i* on patch *l* via emigration (*E_i,l_*) from this patch and immigration (*I_i,l_*) into the patch. The full population dynamics of species *i* on patch *l*, comprising both local, trophic dynamics, Eqs. (2), and dispersal dynamics, can thus be written as

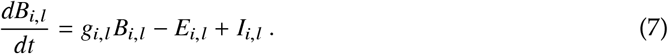

Emigration is a complex process in nature possibly involving different environmental cues and species properties. Here we assume that emigration depends on the net per capita growth rate *g_i,l_* of species *i* on patch *l*, reflecting its current situation in this habitat. If a species’ net growth is positive, there is little need for dispersal and emigration will be low. However, if the local environmental conditions deteriorate, e.g. due to low resource availability or high predation pressure, the emigration rate increases. This is captured by the following function:

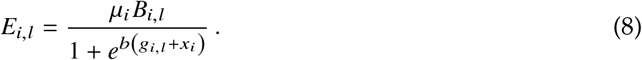

The parameter *μ_i_* = *μ*_0_*x_i_* determines the maximum per capita emigration rate and *b* determines how sensitively the emigration rate depends on the net growth rate (i.e., how quickly it drops when *g_i,l_* increases). Finally, immigration of species *i* into patch *l* depends on the amount of emigration from all neighboring patches *k* as well as on the specific dispersal network, encoded in the success and weight functions *S_i,lk_* and *W_i,lk_*, according to

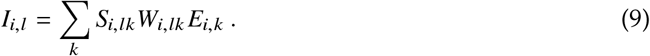

The parameters defining the dispersal dynamics are also summarized in Tab. 1.

### 2.3 Simulation setup

#### 2.3.1 Static and dynamic landscapes

The baseline simulations are carried out using static landscapes, i.e., with RGG networks of *Z* = 30 habitat patches as described above, where all patches and dispersal links are permanently available. However, since the environmental conditions in nature are rarely completely constant, we also study dynamic landscapes in which a fraction *σ* of the patches becomes periodically unavailable as a habitat. This process is called ‘blinking’ and has a period length *λ*. Blinking patches are turned on and off synchronously and change their state every 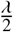 time units. When the blinking patches are turned off, the local food chains go extinct immediately. Furthermore, the dispersal network can be disrupted because these patches cannot be used as stepping stones for dispersal between patches that are too far apart for a direct dispersal link.

#### 2.3.2 Landscape connectance

To capture the effects of varying habitat fragmentation, the intercept of the maximum dispersal distance, *D*_0_, (Eq. (4)) is varied systematically between 0.06 and 0.5. This creates habitat networks that range from mostly isolated patches to systems where the predator can move in a single step between any two patches. The degree of habitat fragmentation is quantified by the connectance of the predator’s dispersal network,

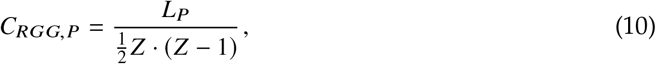

with *L_P_* the number of undirected dispersal links of the predator and Z the number of habitat patches. Note that a higher landscape connectance implies a less fragmented habitat. Also note that using the connectance of the dispersal network of any of the other species to define landscape connectance would only rescale the x-axis of the results (Fig. 3), but not change them qualitatively.

**Fig. 3.**
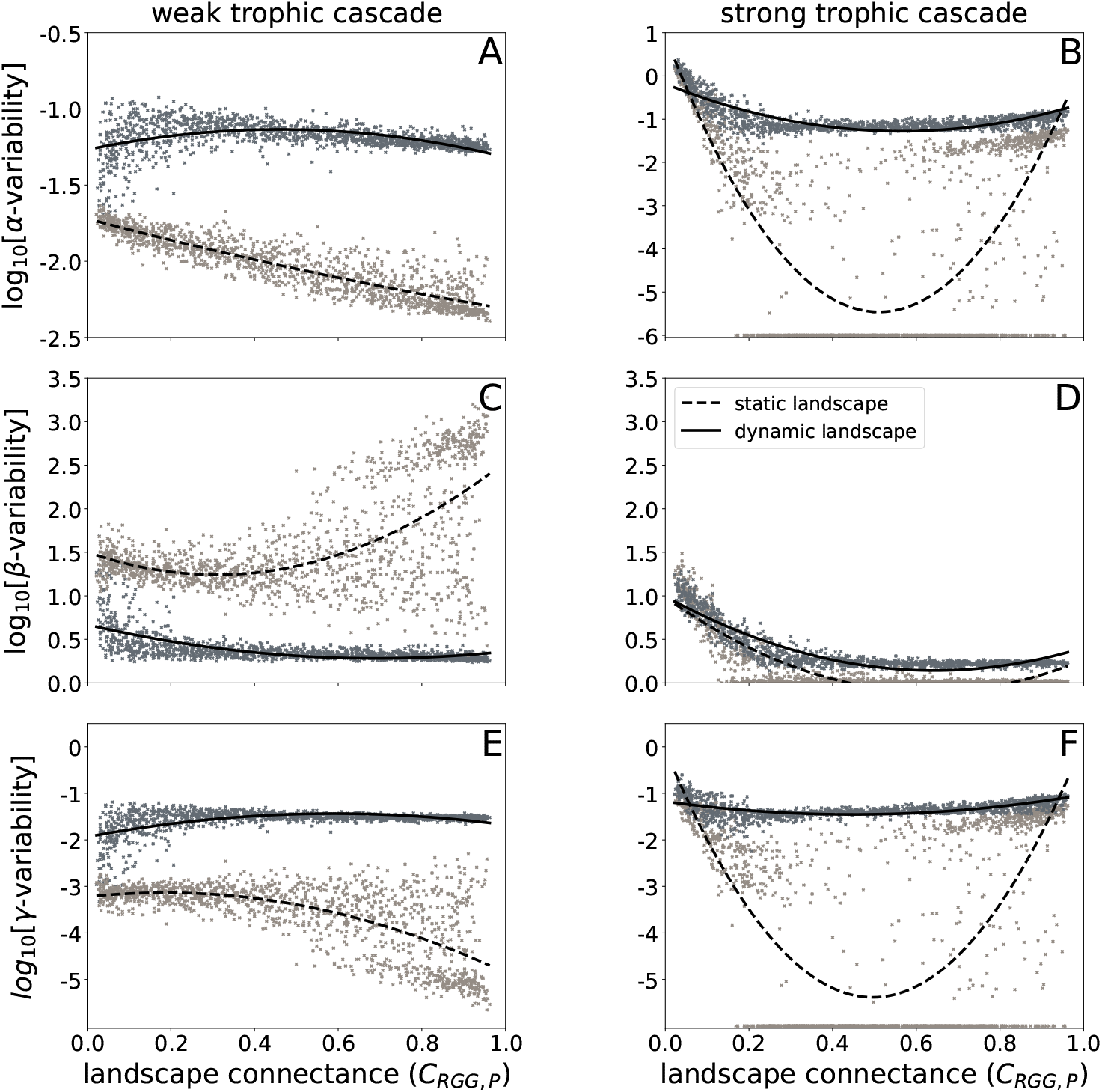
Local (*α*-variability, top row), between patch (*β*-variability, middle row) and metapopulation dynamics (*γ*-variability, bottom row) of the predator for the weak (left column) and the strong trophic cascade (right column). Light grey data points and dashed trend lines (second order fit) indicate static landscapes, dark grey data points and solid trend lines indicate dynamic landscapes. Each data point represents the result of one simulation run with a unique spatial network of habitat patches. All data points where the variability is below 10^−6^ are set to 10^−6^ as differences between them provide no meaningful information that close to the fixed point.

#### 2.3.3 Ecosystem stability

We evaluated ecosystem stability according to Wang and Loreau (2014) as *α*-, *β*-, and *γ*-variability of autotroph, consumer, and predator. For the local or *α*-variability of a species, the coefficients of variation (CV, 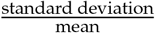) of its local biomass densities on all patches are calculated and then averaged, while for the *γ*-variability (variability of the metapopulation) the CV of the total biomass density (sum over all patches) is evaluated. Similar to the *α*-, *β*-, and *γ*-diversity indices (Whittaker, 1972), *β*-variability measures differences between the patches and can thus be used to determine how synchronously local biomass densities on the different patches oscillate. It is here defined as 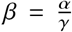. In contrast to the diversity indices, however, variability decreases with an increase of spatial scale, i.e. *γ* ≤ *α* and thus *β* ≥ 1. Spatially synchronous oscillations result in a low *β*-variability and a *γ*-variability that approaches the value of the *α*-variability. Perfect synchronicity is obtained at *β* = 1. An intuitive example of two species, one with synchronous and one with asynchronous oscillations, is provided in the Online Resource (Fig. S1).

#### 2.3.4 Numerical simulations

We simulated food chains that were parameterized to exhibit either a strong or a weak trophic cascade, corresponding to a very uneven or a relatively even distribution of biomass along the food chain, respectively. The spatial networks were either static (all patches permanently available as habitats) or dynamic (30% of the patches periodically becoming unavailable as habitats). The landscape connectance was varied gradually by increasing *D*_0_ from 0.06 to 0.5 in steps of 0.01. Simulations were carried out with a full-factorial design and 30 replicates for each combination of parameters, resulting in a total of 5400 simulation runs. Replicates differed in the randomly chosen positions of 30 patches that formed the spatial networks. Time series were simulated for 90 000 time units and split in three sections of equal length. During the first section, the systems settled on the attractor and from the second section, mean biomass densities were calculated. These mean biomass densities were then used to calculate the variability coefficients from the third section of the time series. During the simulations, a species was considered extinct on a given patch if its local biomass density fell below 10^−20^ and globally extinct if its total biomass density fell below 10^−10^. The local extinction threshold had to be lower than the global threshold to allow for recolonization of blinking patches. Numerical simulations of the ODE model were performed in C (source code adopted from (Schneider et al., 2016)) using the SUNDIALS CVODE solver (Hindmarsh et al., 2005) with absolute and relative error tolerances of 10^−10^. Output data were analysed using Python 2.7.11, 3.6 and several Python packages, in particular NumPy and Matplotlib (Hunter, 2007, Oliphant, 2015, Van der Walt et al., 2011).

## 3 Results

### 3.1 Food chain dynamics without dispersal

To capture how different parameterizations of trophic interactions affect the metacommunity dynamics, we analysed two contrasting trophic cascades in the food chain that were created by assuming either low or high attack rates. The first type, called weak trophic cascade, (*a_CA_* = 105, *a_PC_* = 450, Fig. 2A) is characterised by a weak predation pressure of the predator, a relatively even distribution of biomass along the food chain and a high oscillation frequency (note the different scales of the x-axes of the two panels in Fig. 2). The strong trophic cascade (*a_CA_* = 170, *a_PC_* = 10000, Fig. 2B) is, in contrast, characterised by a very uneven distribution of biomass with a strong dominance of the autotroph (caused by the suppression of the consumer by the predator), a much lower oscillation frequency, and much more drastic population cycles that drive both the predator and the consumer biomass densities repeatedly to very low values. The difference between the predator attack rates in the two cases had to be this pronounced as for intermediate values, the food chain is stable and the analysis of (meta-)population variabilities is not possible (Online Resource, Fig. S2).

### 3.2 Metacommunity dynamics

We evaluated the two different landscape scenarios (static vs. dynamic) for both the weak and strong trophic cascade on a spatial network of *Z* = 30 habitat patches over a gradient of the landscape connectance. All scenarios are evaluated with respect to local (*α*-variability), between patch (*β*-variability), and metapopulation dynamics (*γ*-variability). The observed trends in population variabilities on the different spatial scales were always the same for all trophic levels. We therefore only show results for the predator species. Results for the autotroph and consumer species are in the Online Resource (Figs. S3 and S4).

#### 3.2.1 Local dynamics: α-variability

In contrast to our expectations, a higher landscape connectance dampens biomass oscillations in static landscapes (decreasing *α*-variability, Fig. 3A,B). This trend is particularly pronounced for low to intermediate connectance values in the strong trophic cascade (Fig. 3B), where many systems even settle on a stable fixed point (*α*-variability ≤ 10^−6^). Only at high connectance values we observe a slight increase of the *α*-variability, leading to an overall u-shaped pattern. In the weak trophic cascade, *α*-variability monotonously decreases with landscape connectance, but no stable fixed point is reached.

In dynamic landscapes, *α*-variability is higher than in static landscapes, but it varies only weakly with connectance. In the weak trophic cascade, *α*-variability initially increases, but then decreases slightly, while in the strong trophic cascade it remains largely constant after an initial decrease.

#### 3.2.2 Between patch dynamics: β-variability

On the regional scale we evaluated to what extent the biomass dynamics between habitat patches synchronized (Fig. 3C,D). In line with our expectations, there is in most cases a clear trend towards synchronization of the dynamics by increasing the landscape connectance. Almost perfect synchronization (*β*-variability ≈ 1) is attained for the strong trophic cascade in static landscapes (Fig. 3D), while in the case of dynamic landscapes the level of synchronization seems limited (*β*-variability ≈ 2 for both weak and strong trophic cascades). However, this is only a numerical effect due to the difference between constant and blinking patches.

Only the weak trophic cascade in static landscapes deviates from the general trend: The *β*-variability is not only higher than in the other cases, but it also only slightly decreases from low to medium landscape connectance. For higher connectance the trend levels off and the scatter in the data points increases. Also, for *C_RGG,P_* ≳ 0.6 a separate cloud of data points with very high *β*-variabilities emerges, suggesting that in this part of the parameter space a second attractor with even less synchronization between the patches exists. The bistability of the system is indeed confirmed by dedicated simulations using spatial networks with fixed coordinates of the patches (cf. Online Resource, Fig. S5)

#### 3.2.3 Metapopulation: γ-variability

For both the weak and the strong trophic cascade we find a relatively constant total biomass of the metapopulation (*γ*-variability < 10^−1^, Fig. 3E,F). As expected, *γ*-variability is higher in dynamic landscapes than in static ones. Since local biomass oscillations are often highly synchronized, the trends in the metapopulation dynamics largely follow those already observed in local dynamics: in static landscapes the *γ*-variability declines for the weak trophic cascade and exhibits a valley-shaped pattern for the strong trophic cascade, while in dynamic landscapes it is almost constant in both cases. As with the *β*-variability of the weak trophic cascade in static landscapes, at higher values of the connectance (*C_RGG_* ≳ 0.6) a small cloud of data points appears to be separated from the rest, which have a low *γ*-variability. Again, these data points can be attributed to an alternative attractor with less synchronized dynamics and correspondingly a lower *γ*-variability.

## 4 Discussion

The impact of habitat fragmentation on biodiversity and community dynamics is a subject of on-going debate (Fahrig et al., 2019, Fletcher et al., 2018). Here, we used landscape connectance as a metric of fragmentation and evaluated its effects on the population dynamics of two contrasting states of a meta-food chain (strong and weak trophic cascade). Landscapes were formed by complex networks of habitat patches that were either all permanently available (static landscapes) or partly only seasonally available (dynamic landscapes). By continuously varying the degree of landscape connectance we accounted for potentially interacting effects of temporal and spatial variation of the habitat network. Throughout the discussion we use the term stability in the context of how much temporal variability of biomass is present on the local and the metapopulation level and how synchronized these dynamics are. The discussion focuses on the (meta)population of the predator, but the conclusions also hold for the other two species or the entire metacommunity.

### Main trends

In static landscapes, the local biomass (*α*-) variability decreases with increasing landscape connectance in both the weak and the strong trophic cascade. In the latter case, the dynamics even settle at a fixed point for intermediate values of the landscape connectance, leading to a u-shaped overall pattern. However, regarding the synchronicity of the dynamics (*β*-variability), the two tropic cascades differ significantly: While we observe almost perfectly synchronous dynamics at intermediate to high landscape connectance in the strong trophic cascade, in the weak trophic cascade *β*-variability is much higher (i.e., the dynamics are asynchronous) and even increases with connectance. This increase is mainly driven by the emergence of a second attractor with very asynchronous dynamics at a higher landscape connectance (Fig. 3C and Tab. 2). Regarding the metapopulation (*γ*-) variability, we observe a neutral to decreasing trend in the weak trophic cascade (where the decrease is again driven by the emergence of the second, asynchronous, attractor), while in the strong trophic cascade the pattern is u-shaped, as for the *α*-variability. However, in contrast to the *α*-variability, the *γ*-variability not lower at high landscape connectance than at low landscape connectance. This lack of stabilization of the metapopulation dynamics is caused by the synchronization of the dynamics with increasing landscape connectance.

**Table 2.** Summary of the trends of *α*-, *β*- and *γ*-variability with increasing landscape connectance for the weak (WTC) or strong (STC) trophic cascade in static or dynamic landscapes.

| State of landscape | Type of effect | Trend for WTC | Trend for STC |
| --- | --- | --- | --- |
| static | α-variability | ↓ | u-shape |
| static | β-variability | ↘ & ↑ | ↓ |
| static | γ-variability | → & ↓ | u-shape |
| dynamic | α-variability | → | ↘ |
| dynamic | β-variability | ↘ | ↓ |
| dynamic | γ-variability | ↗ | → |

Dynamic landscapes do not qualitatively change the observed trends in the strong trophic cascade, but significantly weaken them (Tab. 2). Overall, variabilities are higher, and the local and metapopulation dynamics do not settle at a fixed point at intermediate values of landscape connectance anymore. In the weak trophic cascade, dynamic landscapes also increase *α*- and *γ*-variability (which are now also almost independent of the landscape connectance), but the trend of the *β*-variability is reversed and the dynamics almost perfectly synchronize at medium to high landscape connectance. Additionally, in dynamic landscapes no bistability is observed in the weak tropic cascade anymore.

### The role of connectance in static landscapes

Most theoretical studies addressing the stability and possibly synchronization of the population dynamics in metacommunities use only two patches (e.g. Abrams and Ruokolainen (2011), Gramlich et al. (2016), Koelle and Vandermeer (2005)) or regular spatial structures such as lines, rectangular grids, or other spatially symmetric patch arrangements (Plitzko and Drossel, 2015, Sherratt et al., 2000). A notable exception is Holland and Hastings (2008), but the completely random structure of the patch networks analysed there leads to geometrically implausible patch arrangements. In contrast, we study complex networks of habitat patches where the patches still have well-defined geometrical positions. As a result, the model resembles more realistic landscapes and allows us to directly address ecologically relevant processes like habitat fragmentation.

Our theoretical model allows us to study a single aspect of fragmentation, namely landscape connectance (referred to as habitat patch isolation by Fahrig (2003)). The model does not require the definition of an effective patch size, which omits the difficulties of separating the effects of habitat loss from fragmentation within a landscape and also excludes effects commonly accompanying fragmentation, such as edge effects (Pfeifer et al., 2017). Landscape connectance is commonly evaluated in conservation studies (Lindborg and Eriksson, 2004, Urban and Keitt, 2001) and has a significant effect on the dispersal abilities of species. A largely connected landscape allows for higher dispersal rates, which has been shown to synchronize the dynamics of metacommunities (Gouhier et al., 2010), thereby increasing the risk of correlated extinctions throughout the entire landscape (Heino et al., 1997). Our results confirm an increasing synchronization of the dynamics mostly for the strong trophic cascade, but to a lesser extent (from low to intermediate landscape connectance) also for the weak trophic cascade. Contrary to what was observed by Koelle and Vandermeer (2005), there are no opposing trends in synchronisation for the different species along the food chain. However, the stability of the metapopulation (*γ*-variability) is actually to a large extent driven by the response of the *α*-variability to changes in the landscape connectance. For the weak trophic cascade in particular, we observe that – in contrast to our initial hypothesis – *α*-variability continuously declines with increasing landscape connectance, thereby also stabilizing the metapopulation dynamics.

The hypothesis that local variability should increase with landscape connectance was based on the principle of energy flux (Rip and McCann, 2011), according to which decreasing (dispersal) mortality at higher landscape connectance should strengthen and thereby destabilize the trophic interactions along the food chain. In contrast to the prediction of Rip and McCann (2011), low dispersal mortality does not generally result in higher *α*- or even *γ*-variability in our model. We attribute this counter-intuitive trend to an indirect effect of dispersal mortality: Despite their superior dispersal abilities, top predators often suffer most from landscape fragmentation because they are energetically more limited than the species on lower trophic levels (Ryser et al., 2019). In fact, we also find that the lower the landscape connectance, the lower the mean biomass of the predator (see Online Resource, Fig. S6). This decreases the per-capita predation mortality of the consumer, which more than compensates for the increase in the consumer’s dispersal mortality. In line with the principle of energy flux, this destabilizes the consumer-autotroph interaction. At low landscape connectance, the *α*-variability of the predator thus increases because the dynamics of the predator is driven by the increasingly unstable consumer-autotroph interaction.

### Bistability in the weak tropic cascade

In static landscapes, the weak trophic cascade is bistable for medium to large landscape connectance: in addition to the attractor with intermediate (and slightly increasing) synchronicity, which exists for the entire range of landscape connectance, a second attractor with very asynchronous dynamics between the patches emerges. Interestingly, the bistability concerns only the synchronicity of the dynamics (and consequently the *γ*-variability). Local (*α*-) variability is not affected by whether the populations on different patches cycle more or less in synchrony (Fig. 3A).

Bistability in a meta-food chain with three species is already reported by Jansen (1995), who shows the coexistence of a stable limit cycle and a three species equilibrium, when a single species is dispersing. In our model, where all three species are able to disperse, two stable limit cycles are able to exist simultaneously. A possible explanation for the occurrence of the alternative synchronization patterns is the way the dispersal rate is modeled. Specifically, that the rate at which individuals emigrate from a given patch depends on the net growth rate they experience there. Emigration can thus be driven by a lack of resources (in which case emigration helps ending the unfavourable growth conditions and is thus self-limiting) or by an exceedingly high predation rate (in which case emigration actually intensifies the per-capita predation rate for the remaining individuals and becomes self-enforcing). Preliminary analyses suggest that dampening or amplification of net dispersal flows by synchronous and asynchronous oscillations, respectively, create different feedback loops based on these different drivers of emigration, but more detailed analyses are required to understand in detail how these contrasting states stabilize themselves.

### Effect of dynamic landscapes

Seasonality is a nearly ubiquitous feature of ecological systems, since in most environments there are seasonally fluctuating climatic drivers present (Fretwell, 1972). Unlike in other studies where changing environmental conditions modify species traits directly (e.g. respiration rates that depend on temperature), seasonality here affects the habitability of some of the patches. Hence, seasonality periodically changes the connectance of the landscape as perceived by the species. A natural example of such seasonally dynamic landscape are kettle holes which have a species rich community during the colder and wetter season, but run dry during the summer (Kalettka and Rudat, 2006). For simplicity we exclude the occurrence of resting stages in our model, which can play a critical role in the process of recolonization of such habitats (Wade, 1990). This assumption further promotes synchronicity, since an independent restart of the local communities in the blinking patches is not possible.

In our model, dynamic landscapes destabilize the meta-food chain (larger *γ*-variability), which is mostly due to an increase in local (*α*-) variability. This is of course to be expected when an external perturbation is applied: a main driver of the larger local oscillations is the blinking of patches itself, which causes low frequency biomass oscillations through the extinction- and recolonization process and decreases the mean biomass densities on these patches. In the weak trophic cascade seasonality also synchronizes the metacommunity dynamics, which contributes to the strong increase in *γ*-variability in this case. Again, this effect is not unexpected, as environmental noise such as seasonality has long been known to be able to synchronize ecological dynamics in coupled habitats (Moran, 1953). The alternative attractor with very asynchronous dynamics disappears, and for all but the lowest values of landscape connectance almost full synchronization throughout the entire metacommunity is observed. This strong effect is a bit surprising, considering that seasonality in our model constitutes actually only a quite small perturbation. While it periodically leads to the complete extinction of almost a third of the local communities, it does so very rarely: The period length of the full seasonality cycle is *λ* = 6000, which is about 150 times slower than the period length of the population cycles in the weak trophic cascade.

When landscapes are dynamic, weak and strong trophic cascade respond very similarly to landscape fragmentation. Only at very low landscape connectance some asynchronous oscillations can be observed, otherwise the metacommunity dynamics are in both cases almost perfectly synchronized. This indicates that seasonal effects, even if they are relatively weak or act on a much slower time scale than the ecological interactions, have a stronger influence on ecosystem stability than the identity of the species in the community, the strength of the trophic interactions between them, or even the degree to which a landscape is fragmented. This might explain why effects of fragmentation ‘per se’ are often found to be quite small (Fahrig, 2003).

### Relevance and effects of dispersal assumptions

Dispersal comprises a large variety of strategies, and a multitude of causes affects an individual’s decision to leave its home patch (Bowler and Benton, 2005). Causes of dispersal are for example intraspecific competition (Herzig, 1995), quality of food resources (Kuussaari et al., 1996), or top-down pressure through parasitism or predation (Sloggett and Weisser, 2002). In our model we use the net growth rate of a species in a given patch to determine whether the individuals tend to stay or leave this patch. Since the net growth rate depends on both food availability and predation pressure, the model captures multiple of the above mentioned causes of dispersal. However, we assume that individuals have only knowledge about the growth conditions in the patch they are currently in and not about the conditions in potential target patches. The dispersal rate thus only depends on the local conditions and not on the difference in net growth rates between source and target patches. Using a consumer-resource model with two patches, Abrams and Ruokolainen (2011) showed that when the dispersal rate depends on the difference of the growth rates between source and target patch, asynchronous (antiphase) cycles frequently occur, which promotes stability. With our approach, we only find asynchronous dynamics in static landscapes, but even then synchronous metacommunity dynamics frequently occur.

The details of the model for the dispersal rate can have effects beyond the question whether or not the population dynamics on the different patches synchronize. For example, Gramlich et al. (2016) analyzed 19 different dispersal models in a two-patch, two-species system to determine under what conditions population oscillations arise in the first place. They showed that dispersal often destabilizes otherwise stable population dynamics, especially when dispersal rates are constant (random dispersal). With density-dependent (i.e., non-random) dispersal, on the other hand, in some models a positive correlation between dispersal rate and stability emerged. This corresponds to some extent to our findings, where such a stabilization of the dynamics can be seen in the strong trophic cascade at intermediate landscape connectance.

### Conclusion

Our model simulations confirm that a largely connected landscape promotes synchronization of biomass dynamics in a metacommunity (Gouhier et al., 2010), but it also dampens local oscillations. Overall, increasing landscape connectance therefore does not lead to increased biomass variability on the metacommunity level. Without seasonality we even find in the weak trophic cascade an alternative stable limit cycle with asynchronous biomass dynamics. Thus, while dispersal is sometimes considered as a “double-edged sword” (Hudson and Cattadori, 1999), since too much of it can synchronize metacommunity dynamics and thus increase the risk of correlated extinctions, this seems to be less of a problem when dispersal is adaptive, more ecological interactions than just a single trophic link are present, and realistic landscapes consisting of a complex network of habitat patches are considered. However, when accounting for seasonality, the picture reverses again as both local and metacommunity variability increase, and increasing landscape connectance always leads to almost fully synchronized dynamics. Thus, depending on whether environmental conditions are constant or not, the degree of fragmentation of a landscape can have a larger (without seasonality) or smaller impact (with seasonality) on metacommunity stability.

We conclude that in each unique landscape, comprising a multitude of abiotic factors, the impact of seasonally changing environmental conditions has the potential to outweigh other local interactions present in a community. The effect of fragmentation on the stability of a metacommunity thus may strongly depend on local environmental conditions which are relevant for reliable predictions. Furthermore, the non-monotonous stability response curve of the strong trophic cascade shows that the effect of fragmentation on a species or metacommunity may not be trivial and there might be transitions where fragmentation ‘per se’ might switch from having a positive to having a negative effect on the stability of a community.

## Acknowledgement

This study was financed by the German Research Foundation (DFG) in the framework of the research unit FOR 1748 - Network on Networks: The interplay of structure and dynamics in spatial ecological networks (GU 1645/1-1). Further thanks to R. Ceulemans and S. Bolius for critical remarks on the manuscript.

## Author contributions

All authors conceived the study design. MB and MS wrote the computer code. MS performed the numerical simulations and evaluated the data. MS and CG interpreted the results. The first draft of the manuscript was written by MS, editing was lead by CG. All authors read and approved the final manuscript.

## Code availability

We will enable full reproducibility of our study by providing the original C- and Python-code when needed/the manuscript is accepted

