## Supplementary material for "Interaction of habitat isolation and seasonality: impact of fragmentation in dynamic landscapes on local and regional population variability in meta-food chains"

**Theoretical Ecology manuscript No.**
(will be inserted by the editor)

### Supplementary to " Interaction of habitat isolation and seasonality: impact of fragmentation in dynamic landscapes on local and regional population variability in meta-food chains"

**Markus Stark*** · **Moritz Bach** · **Christian Guill**

the date of receipt and acceptance should be inserted later

*corresponding author

Markus Stark

Institute of Biochemistry and Biology, University of Potsdam, Maulbeerallee 2, 14469 Potsdam, Germany

Moritz Bach

Institute of Biochemistry and Biology, University of Potsdam, Maulbeerallee 2, 14469 Potsdam, Germany

Christian Guill

Institute of Biochemistry and Biology, University of Potsdam, Maulbeerallee 2, 14469 Potsdam, Germany

#### S1 Example Variability

In this section, we show illustrative time series of two species on two patches to demonstrate effects of synchronous and asynchronous dynamics on $\alpha$-, $\beta$- and $\gamma$-variability (Fig. S1.1). Frequency and amplitude of the biomass oscillations (and thus also the $\alpha$-variability) are the same for both species. Perfectly asynchronous (antiphase) oscillations of species 1 result in $\beta$-variability $\rightarrow \infty$ and $\gamma$-variability approaching 0. Conversely, perfectly synchronous dynamics of species 2 result in a $\beta$-variability of 1 and $\gamma$-variability equal to the $\alpha$-variability.

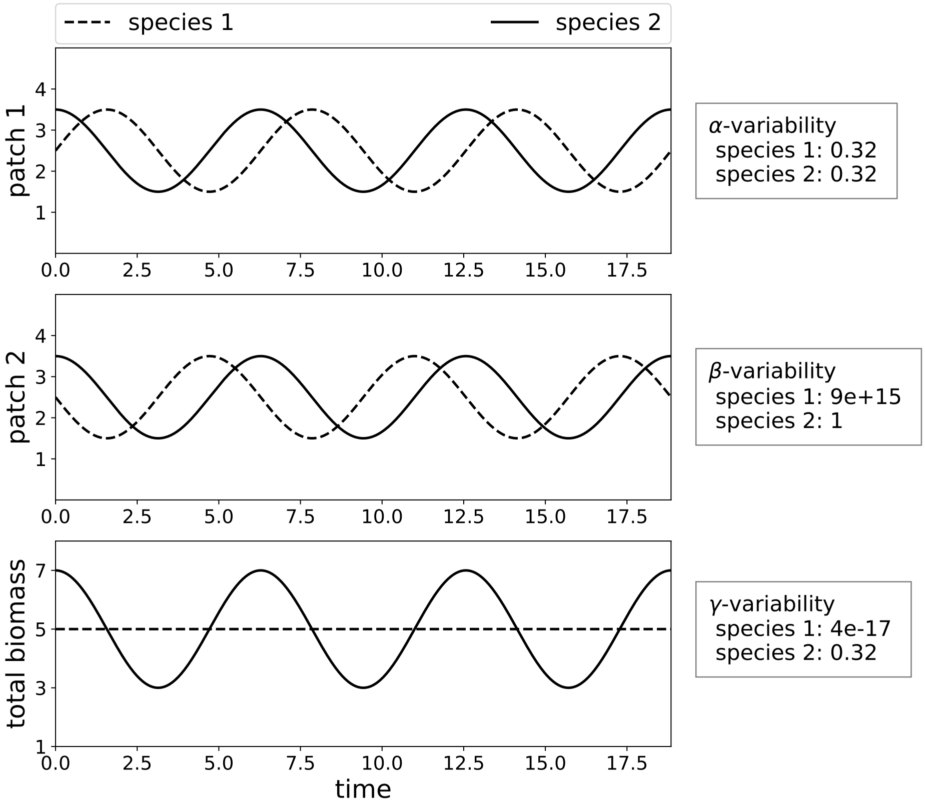


**Fig. S1.1** Exemplary dynamics of a system with two species on two patches. The first two panels show the time series for each species on patch 1 and 2, respectively, while the bottom panel shows the corresponding total biomasses. The boxes to the right contain the numerical values of the $\alpha$-, $\beta$- and $\gamma$-variabilities.

#### S2 Attack rate dependence of local biomass oscillations

We tested a broad range of combinations of consumer and predator attack rates to find suitable parameter sets for oscillatory dynamics. There are two areas in the parameter space that show oscillatory dynamics, separated by an area that leads to a stable equilibrium (Fig. S2.1). Oscillatory behavior is indicated by an increased $\alpha$-variability (CV). Mean biomasses of the species are shown to illustrate the two different trophic cascades. The change in biomass between the two cascades is most striking for the autotroph: at low attack rates (weak trophic cascade), it has relatively low biomass, as it is controlled by the consumer, which in turn is only weakly controlled by the predator, whereas its biomass is high when attack rates are also high (strong trophic cascade). Now the consumer is strongly top-down controlled by the predator and cannot control the autotroph anymore.

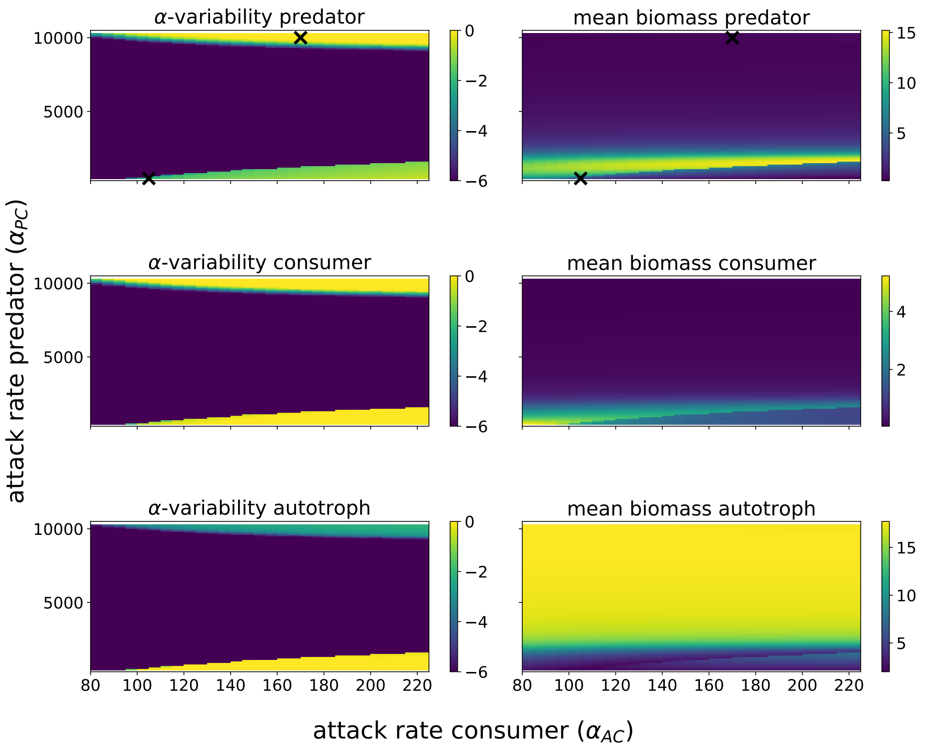


**Fig. S2.1** Simulation results of the food chain on a single patch for a broad range of consumer- and predator attack rates. Shown are the $\alpha$-variabilities (left) on a logarithmic scale ($log_{10}$) and biomasses (right) of predator, consumer, and autotroph species. The black x's denote the parameter values used for the main simulations. Step size attack rate predator: 10; step size attack rate consumer: 5

#### S3 Variabilities of consumer and autotroph species

To complement the results, we here show $\alpha$-, $\beta$-, and $\gamma$-variabilities for the consumer (Fig. S3.1) and the autotroph (Fig. S3.2). The main trends for the local ($\alpha$-variability), between habitat patches ($\beta$-variability) and metapopulation dynamics ($\gamma$-variability) are qualitatively the same for the consumer and the autotroph as for the predator species (Fig. 3).

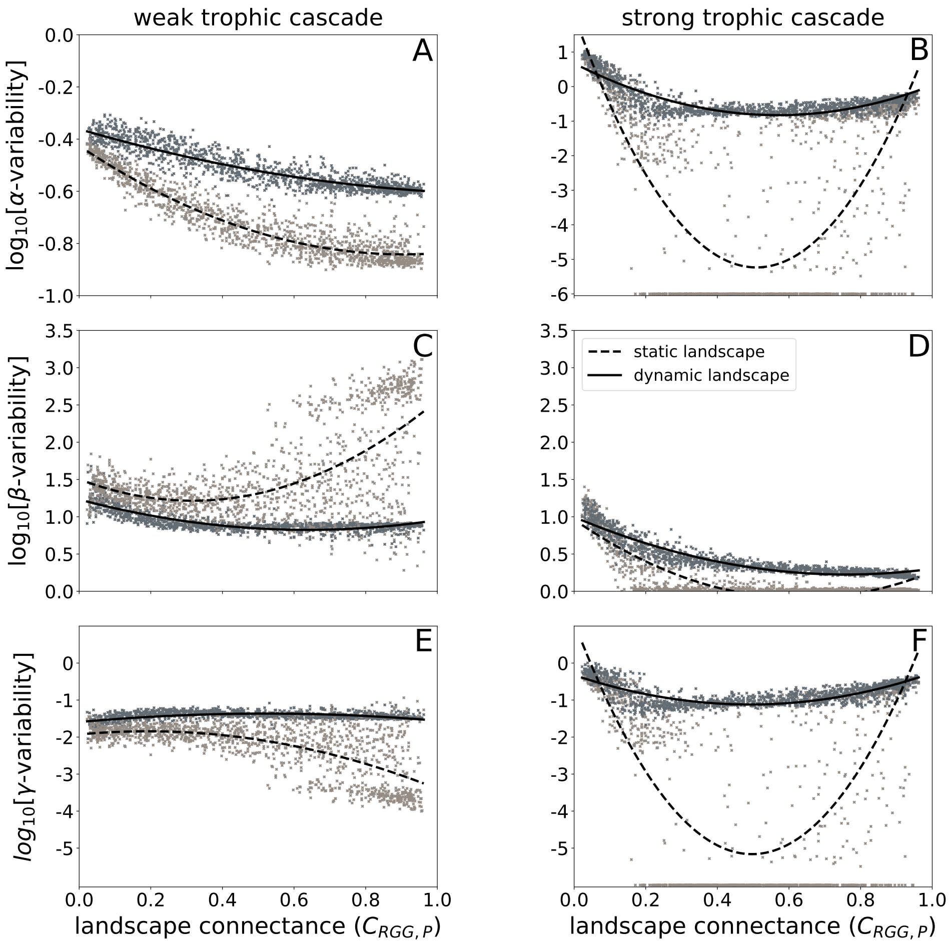


**Fig. S3.1** Local ($\alpha$-variability, top row), between patch ($\beta$-variability, middle row) and metapopulation dynamics ($\gamma$-variability, bottom row) of the consumer for the weak (left column) and the strong trophic cascade (right column). Light grey data points and dashed trend lines (second order fit) indicate static landscapes, dark grey data points and solid trend lines indicate dynamic landscapes. Each data point represents the result of one simulation run with a unique spatial network of habitat patches. All data points where the variability is below $10^{-6}$ are set to $10^{-6}$ as differences between them provide no meaningful information that close to the fixed point.

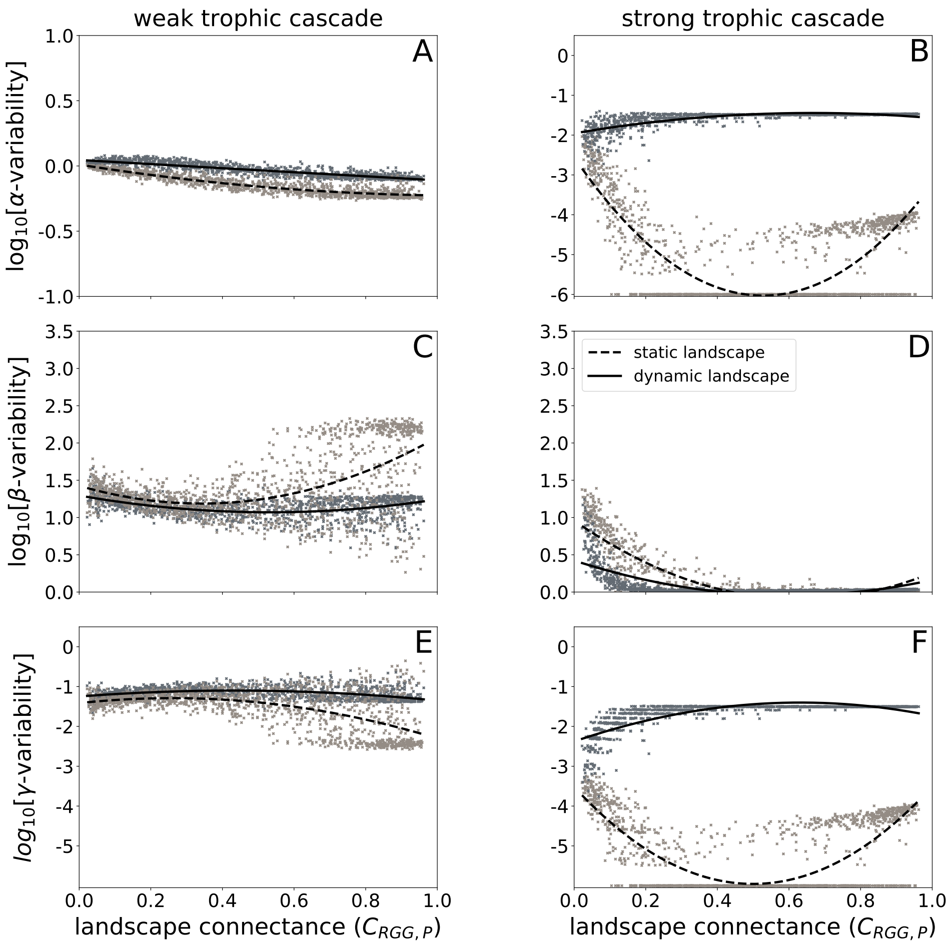


**Fig. S3.2** Local ($\alpha$-variability, top row), between patch ($\beta$-variability, middle row) and metapopulation dynamics ($\gamma$-variability, bottom row) of the autotroph for the weak (left column) and the strong trophic cascade (right column). Light grey data points and dashed trend lines (second order fit) indicate static landscapes, dark grey data points and solid trend lines indicate dynamic landscapes. Each data point represents the result of one simulation run with a unique spatial network of habitat patches. All data points where the variability is below $10^{-6}$ are set to $10^{-6}$ as differences between them provide no meaningful information that close to the fixed point.

#### S4 Alternative stable states in the weak trophic cascade

In order to establish whether the weak trophic cascade (in static landscapes) is indeed bistable, as the seemingly disconnected clouds of data points in Fig. 3 (C and E) suggest, we performed dedicated simulations with one randomly chosen RGG (fixed patch locations) and evaluated the $\beta$-variability of the predator. The minimum dispersal distance, $D_0$, which controls the connectance of the RGG, was varied between 0.06 and 1.05, first from low to high values (grey points in Fig. S4.1) and then from high to low values (black points in Fig. S4.1). In both cases, the respective attractor the system had settled on was numerically followed, but sudden jumps in the $\beta$-variability occurred as the network structure of the RGG changed in a discontinuous way or one of the attractors lost stability. In this particular case, an asynchronous attractor (high $\beta$-variability) is present for all values of $D_0$, while a second attractor with more synchronous dynamics (lower $\beta$-variability) is present only for low ($D_0 < 0.4$) or high ($D_0 > 0.9$) values.

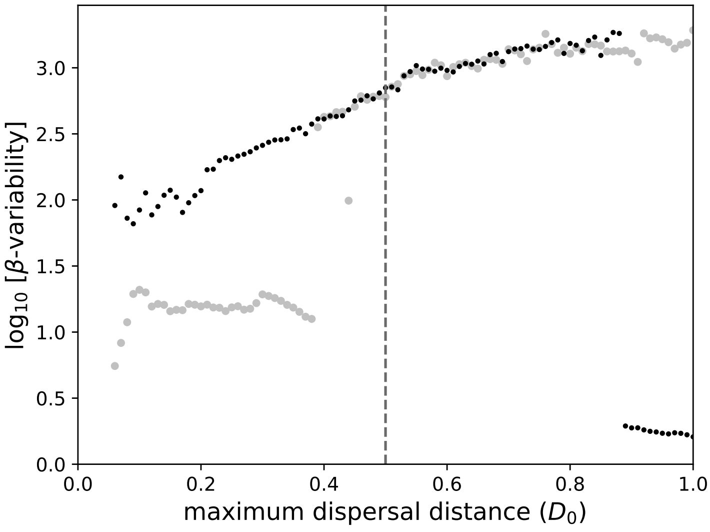


**Fig. S4.1** Bifurcation diagram of the $\beta$-variability of the weak trophic cascade with the minimum dispersal distance, $D_0$, as control parameter. Grey points: Simulations starting at low $D_0$ and gradually increasing it (step size: 0.01), black points: simulations starting at high $D_0$ and gradually decreasing it. The dashed line ($D_0$=0.5) denotes the maximal $D_0$ used for the simulations of the main results.

#### S5 Mean biomasses of each species for different landscape connectances

Over the gradient of landscape connectance, the average biomasses (per patch) of the three species in the weak trophic cascade gradually change. The mean biomass of the predator and consumer species steadily increases with increasing connectance, while the mean biomass of the autotroph decreases.

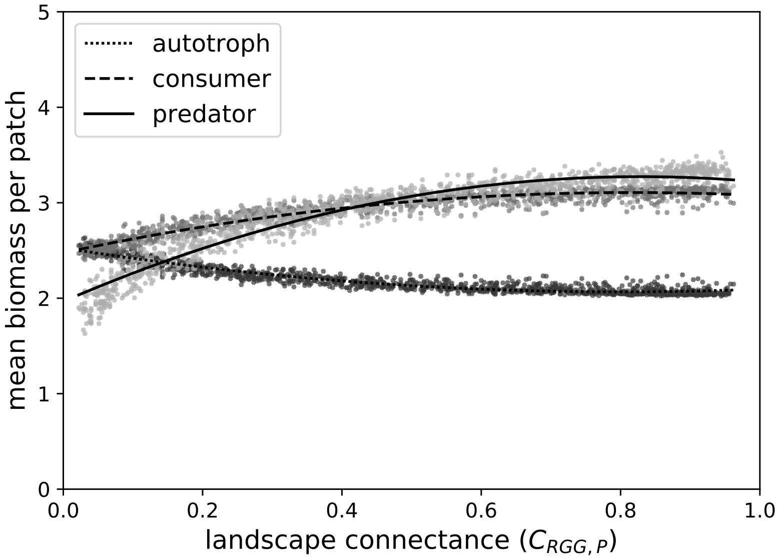


**Fig. S5.1** Mean biomasses (per patch) for the weak trophic cascade in static landscapes. Dotted trend line (second order fit) and dark grey data points: autotroph, dashed trend line and grey data points: consumer, solid trend line and light grey data points: predator. Each data point represents the result of one simulation run with a unique spatial network of habitat patches.
